# Liquid biopsy for infectious diseases: Sequencing of cell-free plasma to detect pathogen DNA in patients with invasive fungal disease

**DOI:** 10.1101/352641

**Authors:** David K. Hong, Timothy A. Blauwkamp, Mickey Kertesz, Sivan Bercovici, Cynthia Truong, Niaz Banaei

**Affiliations:** Karius, Inc, Redwood City, CA, USA; Department of Pathology, Stanford University School of Medicine, Stanford, CA, USA; Division of Infectious Diseases and Geographic Medicine, Department of Medicine, Stanford University School of Medicine, Stanford, CA, USA; Clinical Microbiology Laboratory, Stanford Health Care, Stanford, CA, USA

**Author notes:** Corresponding author: David K. Hong, EMAIL, ADDRESS: 975 Island Drive, Redwood City, CA-94065, Research institution: Karius, Inc., ADDRESS: 975 Island Drive, Redwood City, CA-94065.

**Keywords:** next-generation sequencing, molecular diagnostics, non-invasive, diagnostic biopsy, invasive fungal infection

## Abstract

Diagnosis of life-threatening deep-seated infections currently requires invasive sampling of the infected tissue to provide a microbiologic diagnosis. These procedures can lead to high morbidity in patients and add to healthcare costs. Here we describe a novel next-generation sequencing assay that was used to detect pathogen-derived cell-free DNA in peripheral blood of patients with biopsy-proven invasive fungal infections. The non-invasive nature of this approach could provide rapid, actionable treatment information for invasive fungal infections when a biopsy is not possible.

## 1. Introduction

Invasive fungal infections (IFI) are associated with significant morbidity and mortality in immunocompromised patients, especially those with prolonged neutropenia, allogeneic hematopoietic stem cell transplant (HSCT), solid organ transplants (SOT), prolonged corticosteroids, acquired immunodeficiency syndrome (AIDS), and chronic granulomatous disease (CGD) (Patterson et al., 2016). IFIs result in increased morbidity and mortality for both solid organ and stem-cell transplant patients. Patients with an IFI have 5-fold increased mortality, an additional 19.2 days of hospital stay, and $55,400 in additional costs compared to patients without an IFI (Menzin et al., 2009). Given the high morbidity associated with IFIs, one common approach is to use empiric broad antifungal treatment in patients with persistent fevers after 4-7 days and no identified source, in addition to broad-spectrum antibacterials (Pizzo et al., 1982). The diagnostic evaluation includes extensive imaging to identify a source for the cause of the fever. If a potential deep infection is identified, an invasive biopsy is considered for definitive diagnosis. While this strategy has been employed by many centers, this empiric approach risks overtreatment, with its associated side-effects. Also, invasive biopsies can be costly and have high morbidity. In order to move away from reliance on empiric therapy, management strategies have increasingly relied on non-invasive diagnostic tests to help guide targeted antifungal therapy (Pagano et al., 2010).

The current diagnostic paradigm for invasive fungal disease (IFD) includes categories for proven, probable, and possible disease (De Pauw et al., 2008). These classifications integrate clinical, radiographic, and microbiologic criteria and have been used to guide empiric treatment. Proven IFD requires documented fungal disease by culture, histopathology, or direct microscopic examination of biopsy tissue. However, with the increased usage of serum antigen biomarkers such as *Aspergillus* galactomannan, many patients fall into the probable category, which is defined by clinical findings and additional laboratory evidence. While this provides some directional suggestions that an invasive mold is present, it does not provide species-level identification of the organism which can be critical for guiding therapy. In addition, both the *Aspergillus* galactomannan and beta-D-glucan tests have inadequate sensitivity for *non-Aspergillus* molds including *Rhizopus, Mucor, Fusarium*, and *Scedosporium* (Lamoth et al., 2012) which are increasingly important causes of IFIs (Douglas et al., 2016, Petrikkos et al., 2012). Given the limitations of current diagnostic technology, there is a need for timely, non-invasive, unbiased methods to detect invasive fungal infections particularly in immunocompromised patients (Fishman, 2007, Tomblyn et al., 2009).

We have employed a technique that uses open-ended next-generation sequencing (NGS) of cell-free plasma to detect pathogen DNA (Abril et al., 2016, De Vlaminck et al., 2015). Cell-free DNA (cfDNA) are small fragments of circulating DNA in the cell-free compartment of blood (Mandel and Metais, 1948). NGS and analysis of these DNA fragments, which are derived from dead and dying cells, has been an active area of research and development in many fields of medicine. For example, NGS of cfDNA has been used to diagnose fetal chromosomal defects by detecting fetal cell-free DNA in maternal circulation (Fan et al., 2008). Similarly, NGS has been used to monitor solid organ rejection and tumors by detecting organ-or tumor-derived cell-free DNA in blood (Snyder et al., 2011). Given that the estimated half-life of cfDNA in circulation is less than two hours (Diehl et al., 2008), this approach is uniquely suited to diagnose active infections where there are ongoing dynamics between the pathogen and the immune system. Here we describe a novel application of NGS for non-invasive detection of pathogen-derived cell-free plasma DNA in patients with invasive fungal infections.

## 2. Methods

### 2.1 Ethics

This study was approved by the Stanford University Institutional Review Board (IRB). Discarded plasma samples were collected under a waiver of the informed consent requirement as allowed by the IRB.

### 2.2 Patient samples

Plasma was collected at Stanford University hospital between 3-22-2015 and 8-7-2015. Patient infection was confirmed either by standard fungal culture-based methods or targeted ribosomal locus sequencing for fungi (Gomez et al., 2017). After confirmation of infection, surplus EDTA plasma from clinical testing (complete blood count) was retrieved from the clinical laboratory. Samples were centrifuged, aliquoted, and stored at −80°C for future analysis. Samples and clinical data including confirmed infections were sent to Karius, Inc (Redwood City, CA).

### 2.3 Sample preparation and next-generation sequencing

Plasma samples were thawed, centrifuged at 16,000 rcf for 10 minutes, and spiked with a known concentration of synthetic DNA molecules for quality control purposes. Cell-free DNA was extracted from 0.5 mL plasma using a magnetic bead-based method (Omega Biotek, Norcross, GA). DNA libraries for sequencing were constructed using a modified Ovation^®^ Ultralow System V2 library preparation kit (NuGEN, San Carlos, CA). Negative controls (buffer only instead of plasma) and positive controls (healthy plasma spiked with a known mixture of microbial DNA fragments) were processed alongside patient samples in every batch to ensure that no sporadic environmental contamination was introduced into the assay. Samples were multiplexed with other samples and sequenced on an Illumina NextSeq^®^ 500. On average 45 million reads were obtained for each sample.

#### Analysis Pipeline

Primary sequencing output files were processed using bcl2fastq (v2.17.1.14) to generate the demultiplexed sequencing reads files. Reads were filtered based on sequencing quality and trimmed based on partial or full adapter sequence. The bowtie2 (version 2.2.4) method was used to align the remaining reads against Karius’ human and synthetic-molecules references. Sequencing reads that exhibited strong alignment against the human references or the synthetic molecule references were collected and filtered out from further analysis. The remaining reads were aligned against Karius’ proprietary micro-organism reference database using NCBI-blast (version 2.2.30). A mixture model was used to assign a likelihood to the complete collection of sequencing reads that includes the read sequence probabilities and the (unknown) abundances of each taxon in the sample. An expectation-maximization algorithm was applied to compute the maximum likelihood estimate of each taxon abundance. To determine whether the levels observed in the samples exceeded those expected to originate from the environment alone, a Poisson model parameterized by the estimated background abundances was applied. Only taxa that rejected this null hypothesis at high significance levels were reported and included in downstream analyses. The entire process from DNA extraction through analysis was typically completed within 28 hours. Assuming that shipping from a hospital can take up to 24 hours, results would be available to an ordering clinician in 2-3 days.

## 3. Results

Nine subjects were identified with proven invasive fungal infection. Patients had documented fungal infections from a variety of sources including lung, mediastinal lymph node, heart, brain, sternum, and small bowel (Table 1). Fungal infection was identified either by fungal culture or targeted fungal ribosomal locus sequencing (Gomez et al., 2017).

**Table 1.** Clinical details of patients with invasive fungal infection

| Patient description | Conventional diagnostic procedure | Aspergillus galactomannan | Microbiology result (method) | Plasma NGS results | Plasma collection relative to biopsy | Antifungals at time of plasma collection (days since initiation) |
| --- | --- | --- | --- | --- | --- | --- |
| 73 Y female with multiple myeloma s/p autologous SCT<sup>2</sup> admitted with febrile neutropenia and necrotic bowel | Small bowel resection | Positive (0.73) | *Aspergillus fumigatus* sp. complex by targeted fungal sequencing<sup>1</sup> | *Mycobacterium abscessus*, *Staphylococcus epidermidis* | 20 days after | Caspofungin (18 days)<br>Liposomal amphotericin (11 days) |
| 68 Y male with mixed phenotype leukemia s/p allogeneic SCT<sup>2</sup> presents with right pleuritic chest pain | Lung biopsy | Negative | *Cunninghamella* sp. by targeted fungal sequencing<sup>1</sup> | *Cunninghamella bertholletiae* | 3 days prior | Voriconazole prophylaxis |
| 29 Y woman with SLE<sup>3</sup> admitted with septic shock and respiratory failure | Endotracheal aspirate | Positive (>2.0) | *Aspergillus fumigatus* sp. complex by culture (50-100 colonies), *Emmericella nidulans* by culture (6 colonies) | *Aspergillus lentulus*, CMV | 10 days after | None |
| 39 Y male with adenoid cystic carcinoma Stage III admitted with bacteremia and pulmonary nodules | Lung biopsy | Positive (>1.01) | *Aspergillus fumigatus* by culture (50-100 colonies) | Negative | Same day | None |
| 20 Y female with aortic valve replacement and aortic root enlargement complicated by mediastinal bleeding requiring ECMO<sup>4</sup> | Sternal tissue biopsy | Not done | *Rhizopus* sp. by culture (7 colonies) | *Rhizopus microsporus*, HSV1, *Staphylococcus epidermidis* | Same day | None |
| 59 Y male heart transplant recipient with pancreatic mass | Pancreatic and peri-pancreatic lymph node biopsy | Positive (>2.0) | *Aspergillus terreus* (4 colonies), *Candida albicans* (<10 colonies), *Pseudomonas aeruginosa* (4+), *Streptococcus* sp. (3+) by culture | *Aspergillus terreus*, *Pseudomonas aeruginosa* | 2 days after | Anidulafungin (1 day) |
| 49 Y female heart transplant recipient admitted with altered mental status and a brain abscess | Brain biopsy | Negative | *Scedosporium/pseudoallescheria boydii* complex (>100 colonies) by fungal culture | *Scedosporium apiospermum*, *Pseudomonas pseudoalcaligenes*, *Mycobacterium mageritense*, Torque tenovirus | Same day | None |
| 50 Y male with Burkitt's lymphoma with right sided facial swelling and orbital cellulitis | Sinus biopsy | Not done | *Rhizopus oryzae* by targeted fungal sequencing<sup>1</sup> | *Rhizopus delemar*, CMV | Same day | None |
| 65 Y male s/p aortic valve replacement with fever of unknown origin and pseudoaneurysm near aortic root | Aortic tissue biopsy | Not done | *Rhizopus* sp. by fungal culture | *Rhizopus microsporus* | Same day | None |

<sup>1</sup>Targeted ribosomal sequencing described in Gomez et. al (2017), <sup>2</sup>SCT: Stem cell transplant, <sup>3</sup>SLE: Systemic Lupus Erythematosus, <sup>4</sup>ECMO: Extracorporeal membrane oxygenation

In seven out of nine cases, plasma NGS testing detected the same fungus identified from the biopsy tissue at the genus level. In some cases, conventional microbiology identified molds only to the genus level (e.g. *Rhizopus* species). The fungi identified by plasma NGS included *Aspergillus terreus* and non-*Aspergillus* molds including *Scedosporium, Rhizopus*, and *Cunninghamella*.

In one case, plasma NGS identified *Aspergillus lentulus* while conventional microbiology identified the organism as *Aspergillus fumigatus* species complex. *A. lentulus* is morphologically identical to *A. fumigatus*, shares 91% sequence homology, and both belong to the *Aspergillus fumigatus* species complex with *A. lentulus* showing decreased susceptibility to many azoles (Balajee et al., 2005). This instance highlights the importance of species-level identification of invasive fungal infections.

In cases where plasma NGS testing did not identify the causal IFI organism, this could be due to the delayed timing of plasma sample collection relative to the biopsy procedure. In one case, the plasma sample was obtained 20 days after the biopsy procedure, after at least 15 days of anti-Aspergillus therapy. While the kinetics of fungal DNA in plasma have not been definitively established, there are data from blood *Aspergillus* PCR indicating that the signal may wane >7 days of treatment (Imbert et al., 2016). In the other negative case, *Aspergillus fumigatus* sequences were present in the raw data but they numbered below the read threshold required for a positive test result. In both cases, *Aspergillus* galactomannan tests were positive at the approximate time of plasma sampling.

In six of nine patients, multiple organisms were identified by plasma NGS. This included detection of cytomegalovirus, human herpes simplex virus-1, and torque tenovirus, a likely non-pathogenic virus that is present with increasing immunocompromised states (De Vlaminck et al., 2013). Bacteria were also detected in four patients including *Staphylococcus epidermidis*, *Pseudomonas aeruginosa*, *Mycobacterium mageritense*, and *Mycobacterium abscessus*. The presence of these organisms could not be confirmed as the cultures sent did not reveal these organisms and/or the appropriate culture was not sent (e.g. AFB culture) although both Mycobacteria are rapidly growing and can grow on routine media.

## 4. Discussion

In this study, we demonstrate the use of sequencing of cell-free plasma to non-invasively detect fungal DNA in patients with proven invasive fungal infection. This technique identified molds at the species level from a variety of body locations showing the ability of cell-free plasma to integrate information from many organs. This approach, in conjunction with radiographic data and other clinical data, can help target antifungal therapy without the need for an invasive biopsy procedure and its associated morbidity. Furthermore, increased breadth of detection allows for identification of a wide range of pathogens including both *Aspergillus* and non-*Aspergillus* molds. Of note, this open-ended approach can also potentially identify organisms that can mimic IFIs such as *Nocardia spp* and *Toxoplasma gondii*.

Despite the availability of a number of new antifungal therapies, invasive fungal infections remain a major cause of morbidity and mortality in immunocompromised patients. This is likely due to both an increase in the intensity of immunosuppressive regimens being used, as well as the widespread use of *Aspergillus-active* antifungal prophylaxis which has been associated with a both a rise in antifungal-resistant organisms (Perlin et al., 2017) and an increase in IFIs due to non-Aspergillus molds such as the Zygomycetes (Ito et al., 2010). These molds, such as *Fusarium* spp, *Scedosporium* spp, and the Mucorales are commonly multidrug resistant making precise identification imperative. Given the wide diversity of pathogenic fungi, there is a critical need for rapid, non-invasive detection of species-level identification of these invasive infections to help guide specific antifungal therapy.

Non-invasive biomarkers for IFI including the *Aspergillus* galactomannan and beta-D-glucan tests have been accepted as adjunctive diagnostic tests by the EORTC/MSG (European Organization for Research and Treatment of Cancer/Invasive Fungal Infections Cooperative Group and the National Institute of Allergy and Infectious Diseases Mycoses Study Group) in the management of IFIs (De Pauw et al., 2008). Although galactomannan has a reported sensitivity of around 70% in hematologic malignancy and stem-cell transplant patients, its performance in other immunocompromised patients such as solid-organ transplant or primary immunodeficiency patients is much worse (Patterson et al., 2016). The major limitation of the use of galactomannan and beta-D glucan is their inability to detect the wide range of fungi that can cause IFIs; including *Rhizopus, Mucor, Fusarium*, and *Scedosporium.* Anti-fungal treatment or prophylaxis can also decrease the level of circulating galactomannan (Marr et al., 2004). Fungal DNA levels in plasma may also be affected by antifungal treatment. Studies of *Aspergillus* PCR have demonstrated that *Aspergillus* DNA in serum remains relatively stable for the first week after treatment, and clearance by 2-3 weeks may be associated with favorable treatment outcomes (Imbert et al., 2016). Furthermore, in a study of urine cfDNA for tuberculosis, pathogen cfDNA increased with treatment (Labugger I et al., 2017). Further investigation will be needed to understand the potentially unique biologic variances of different organisms and to better understand the effects of antifungal treatment on the dynamics of pathogen cell-free DNA to interpret the performance of plasma NGS in patients with invasive fungal infections.

The potential of NGS for infectious disease diagnosis has been demonstrated in recent publications, including employing NGS on cerebrospinal fluid to diagnose neuroleptospirosis (Wilson et al., 2014), and direct sequencing of brain tissue to detect potential pathogens in culture-negative neuroinflammatory disorders (Salzberg et al., 2016). While these applications are promising, they still require an invasive procedure to obtain tissue for diagnosis. This is the first report of a plasma NGS test that can non-invasively identify the causal pathogen in proven IFI. This test may allow for earlier targeting of antifungal therapy and forestall an invasive biopsy, particularly when such a biopsy is delayed or could adversely affect the patient’s condition. Since fungal genetic sequences are identified, it may be possible in the future to identify antifungal resistance genes to help further guide therapy.

## 5. Conclusions

We describe a novel application of next-generation sequencing of cell-free plasma to detect fungal DNA from deep infection sites. This test was able to detect both *Aspergillus* and *non-Aspergillus* molds in patients with proven invasive fungal infection diagnosed with invasive sampling of infected tissue. The use of this non-invasive plasma assay can aid in the diagnosis of deep infections, particularly when an invasive diagnostic procedure is not possible, or in immunocompromised patients where the number of potential pathogens is very large.

## 6. Acknowledgements

This work was supported by Karius, Inc., Redwood City, CA.

